# Non-mycorrhizal root associated fungi of a tropical montane forest are relatively robust to the long-term addition of moderate rates of nitrogen and phosphorus

**DOI:** 10.1101/2021.08.11.455858

**Authors:** Juan F. Dueñas, Stefan Hempel, Jürgen Homeier, Juan Pablo Suárez, Matthias C. Rillig, Tessa Camenzind

## Abstract

Andean forests are biodiversity hotspots and globally important carbon (C) repositories. This status might be at risk due to increasing rates of atmospheric nutrient deposition. As fungal communities are key in the recirculation of soil nutrients, assessing their responses to soil eutrophication can help establish a link between microbial biodiversity and the sustainability of the C sink status of this region. Beyond mycorrhizal fungi, which have been studied more frequently, a wide range of other fungi associate with the fine root fraction of trees. Monitoring these communities can offer insights into how communities composed of both facultative and obligate root associated fungi are responding to soil eutrophication.

Here we document the response of non-mycorrhizal root associated fungal (RAF) communities to a long-term nutrient manipulation experiment. The stand level fine root fraction of an old growth tropical montane forest was sampled after seven years of nitrogen (N) and phosphorus (P) additions. RAF communities were characterized by a deep sequencing approach. As per the resource imbalance model, we expected that asymmetries in the availability of C, N and P elicited by fertilization will lead to mean richness reductions and alterations of the community structure.

We recovered moderately diverse fungal assemblages composed by sequence variants classified within a wide set of trophic guilds. While mean richness remained stable, community composition shifted, particularly among Ascomycota and after the addition of P. Fertilization factors, however, only accounted for a minor proportion of the variance in community composition. These findings suggest that, unlike mycorrhizal fungi, RAF communities are less sensitive to shifts in soil nutrient availability. A plausible explanation is that non-mycorrhizal RAF have fundamentally different nutrient acquisition and life history traits, thus allowing them greater stoichiometric plasticity and an array of functional acclimation responses that collectively express as subtle shifts in community level attributes.

## Introduction

Tropical montane forests are increasingly recognized as important global carbon (C) sinks (Duque et al., 2021; Spracklen & Righelato, 2014). Although the question remains as to how such C is partitioned above- and belowground, available estimates suggest the belowground portion is large (de la Cruz-Amo et al., 2020; Girardin et al., 2013; Moser et al., 2011). In contrast to the tight nutrient cycling observed in lowland tropical rainforests, C and other elements tend to accumulate in thick layers of organic matter along mountainous slopes (Wilcke et al., 2002). While it is unclear what drives such accumulation, it is often attributed to reduced soil biological activity due to cooler temperature regimes (de la Cruz-Amo et al., 2020), water saturation and a topographically induced gradients of nutrient limitation (Werner & Homeier, 2015). As soil nutrient availability, temperature and dryness are likely to increase in tropical mountains due to human induced global change (Cusack et al., 2016; Velescu et al., 2016; Wilcke et al., 2019), there is growing concern about the fate of these soil nutrient stocks. Stated succinctly, the question arises if global change drivers could turn tropical montane forests from sinks into sources of greenhouse gases.

Evidence that soil microbial communities are fundamental for the recirculation of nutrients is accumulating (Delgado-Baquerizo et al., 2020; Wagg et al., 2021). Thus, one of the primary mechanisms by which atmospheric nutrient deposition could affect the emission or stabilization of soil C is by affecting soil microbes. Yet, how exactly microbial diversity or composition links with nutrient cycles is still unresolved (Fierer, 2017; Glassman et al., 2018). One of the most salient gaps in our understanding of such link is that most of the studies that inform this link were conducted in temperate and boreal biomes, which are severely N limited (Fayle et al., 2015; Guerra et al., 2020). Unlike higher latitude forests though, soils in montane tropical forests are moderately N-limited, while sometimes suffering from less severe phosphorus (P) depletion relative to lowland tropical forests (Camenzind et al., 2018; Du et al., 2020). Perhaps because of these fundamental differences, experimental addition of N in tropical forests has led to variety of responses that range from positive to negative, depending on the process under examination and the elevation at which the forest is located (Cusack et al., 2016). By contrast, P addition has led to a consistent increase in the rates of N fixation and decomposition regardless of elevation (Camenzind et al., 2018). Given these discrepancies, it is relevant to investigate how tropical soil microbial communities respond to soil eutrophication, by extending our analysis to the broadest set of environmental conditions in which forests occur.

Fungi are at the base of the soil food web and, together with bacteria, are thought to be largely responsible for the cycling of nutrients in the soil (Fierer, 2017; van der Heijden et al., 2008). Fungi represent the dominant microorganism in the upper organic layer of tropical montane forests in terms of biomass (Cusack et al., 2011; Krashevska et al., 2010) and are also believed to be highly diverse in the tropics (Meiser et al., 2014; Tedersoo et al., 2014; Zhou et al., 2016). Despite such prominence, research on the effects of nutrient availability on tropical fungal communities is rare and focuses heavily on specialized groups within the kingdom (e.g. mycorrhizal fungi, reviewed in Cusack et al., 2016). Beyond mycorrhizal fungi, it is possible that other fungal groups are as important to determine the ecosystem response to soil eutrophication. For instance, recent model estimates indicate that saprobic fungi account for 35% of soil respiration, in contrast to the 5% represented by mycorrhizal fungi (Fatichi et al., 2019). Given most fungal endophytes are thought to associate with the plant via recruitment form the soil (Hardoim et al., 2015; Rodriguez et al., 2009), monitoring how non-mycorrhizal root associated fungal (RAF) communities respond to N and P fertilization can shed light on how soil fungal communities are responding. In addition to this, monitoring RAF communities can offer insights into the occurrence and response of fungal pathogenic or mutualistic guilds, which are highly relevant to ensure the co-occurrence of a diverse tree community (Bachelot et al., 2015). Thus by targeting RAF communities, it is possible to increase our understanding of the response of several often neglected components of fungal communities to soil eutrophication.

Predicting the consequences of increased nutrient availability for RAF communities is complex due to our incomplete understanding of how RAF diversity and community structure relate to soil nutrient availability. Notwithstanding this situation, we can employ the existing ecological theory as a predictive framework. The resource ratio model postulates the balance of resources will determine whether co-existence is possible between a set of competing species (Cardinale et al., 2009). If nutrient addition leads to resource imbalances between the substrate and organisms (Li et al., 2016; Peñuelas et al., 2013), and such imbalances are impossible to bridge, differential abilities to obtain the most limiting resource could lead to extinctions. Short term fertilization with N and P in tropical montane forests has favored a greater N and P return to the soil as leaf litter, and an increase in the production of fine root necromass after the addition of P (Homeier et al., 2012).

Together, these responses might affect the composition of C pools belowground. That is, an increased production of leaves and fine roots with shorter lifespans could signal an increased proportion of recalcitrant C inputs, yet a reduction in the exudation of more labile C compounds from living roots. Alterations of the C pool composition, and the N:P ratio of the substrate are key, given fungal stoichiometry appears to closely track the stoichiometry of soils in montane tropical forests (Nottingham et al., 2015). Indeed, non-mycorrhizal soil fungal communities have increased in biomass after the simultaneous addition of a C source together with a mineral nutrient in these forests (Krashevska et al., 2010; Nottingham, Hicks, et al., 2018). Hence, it is reasonable to expect that asymmetric rates of nutrient addition will elicit the response of RAF communities. The direction and magnitude of such change, however, is still unclear.

To test these ideas, we surveyed the fine root fraction of an old growth tropical Andean forest subjected to long-term nutrient manipulation (Homeier et al., 2012). We characterized the taxonomic and trophic guild structure of RAF communities by means of a meta-barcoding approach, and estimated community level attributes across fertilization treatments. To further our assessments of the effects of fertilization on RAF communities, we compared the relative read abundance of higher level taxonomic and guild categories across treatments with a differential abundance method that corrects for distortions in these parameters. We expected that an increase in the availability of organic and inorganic N and P in the soil will lead to 1) a reduction in RAF community diversity because fungal taxa for which the altered resource availability is detrimental will be suppressed; 2) shifts in community composition, not only due to reductions in diversity, but also due to the colonization of empty niches or the competitive dominance of taxa with better dispersal abilities or more flexible coping mechanisms; and 3) major fungal lineages or trophic guilds will respond differently to fertilization in terms of adjusted relative read abundances, due to evolutionary fixed differences in their abilities to obtain limiting resources.

## Materials and Methods

### Experimental design and data collection

Samples were collected in August 2015, from a full factorial nutrient manipulation experiment established in 2007 (NUMEX, Homeier et al., 2012). Experimental plots are deployed in an old growth forest located at ∼ 2100 m. a. s. l. along the southern slopes of the San Francisco river valley – an eastern Andean valley located in southern Ecuador. The experiment lies within San Francisco biological reserve (3°58’S, 79°04’W), which borders with Podocarpus national Park. Based on historical estimates of aerosol deposition (Velescu et al., 2016), the primary goal of NUMEX is to evaluate the ecosystem response to moderate increases in nitrogen and phosphorus deposition. Every six months, 5 kg ha^-1^ of P (as monosodium dihydrogen phosphate; NaH2PO4) and 25 kg ha^-1^ of N (as urea; CH4N2O) have been manually applied to the forest floor. The experiment is organized as a fully randomized factorial block design (Fig. S1). Each block consists of four 400 m2 (20 x 20 m) fertilized plots (i.e. +N, +P, +N+P) and one control plot of equal dimensions.

Within every experimental plot, six cores (40 cm depth and 3,5 cm diameter) distributed along two randomly placed orthogonal transects were collected. The upper 40 cm of soil where NUMEX is established correspond to a nutrient poor organic layer, which is also where the highest fine root density can be found (Moser et al., 2011; Wolf et al., 2011). Upon collection, twenty fine root pieces of approximately 2 cm length and < 2 mm diameter were separated from the upper 10 cm of the organic layer present in each core. Care was taken to maximize the morphological diversity of each mixed root subsample in the hope that this reflects the taxonomic diversity of trees present in the experimental plots (∼45 species, Baez & Homeier, 2018). Upon collection, samples were brought back to the San Francisco research station where roots were rinsed in sterile water, preserved in 97% ethanol and stored at 4° C. Samples were transported to the Institute of Biology of Freie Universität Berlin, were they were stored at −20° C.

### DNA extraction, amplification and sequencing

Isolation of DNA from mixed root samples followed the protocols delineated in Dueñas et al. (2020). Briefly, mixed roots samples were lyophilized and milled. DNA was isolated from the milled material using the MoBio soil kit following the manufacturer’s protocol (MoBio Laboratories Inc., Carlsband, CA, USA).

Primer set fITS7 and ITS4 (Ihrmark et al., 2012) was employed to characterize the variability of the internal transcriber spacer (ITS2) within the fungal rDNA operon. ITS2 amplification and sequencing were obtained from a 50 µl aliquot of each extract (n = 96) that was shipped to Macrogen Inc. laboratories in Seoul, Republic of Korea. Amplification products of the targeted lengths were gel purified and then denatured and ligated to a 5’–3’ adaptor sequence. Libraries were then multiplexed and submitted to the sequencing platform (Illumina MiSeq, pair end 2 x 250 bp). Reads with less than 36 bp were discarded and adaptor sequences were removed by Macrogen Inc.

### Bioinformatics and variant classification

Files of the two sequencing rounds were independently processed with packages Dada2 (Callahan et al., 2016), ShortRead (Morgan et al., 2009) and Biostrings (Pagès et al., 2021) in order to define amplicon sequence variants (ASVs). Both workflows were based on Dada2 developer’s tutorial for ITS sequences (v1.8).

Briefly, primer oligos were removed from every sequence using Cutadap (v.2.1, Martin, 2011). Read ends were truncated if the quality score of base calls fell below two; and filtered out when reads were smaller than 50 bp or presented more than two erroneous or undefined base calls per sequence. Remaining reads were de-replicated and then subjected to the Dada2 sample inference algorithm. Denoised forward and reverse reads were then merged if they had an overlap of at least 12 bp between them. At the end of this process, an ASV abundance table was generated followed by a chimera removal step. Hereafter, the acronym ASV and the term ‘variant’ will be used interchangeably.

To allow comparisons to prior studies, ASVs were clustered into 97% OTUs using functions within the package DECIPHER (Wright, 2020). For all other analysis, ASVs were chosen as taxonomic units over traditionally employed OTUs because: 1) ASVs circumvent the need to specify a fixed similarity threshold to cluster units across fungal lineages with widely differing levels of intra-specific variability (Nilsson et al., 2008); 2) ASVs are directly reproducible and independent of the dataset (Callahan et al., 2017); and 3) ASVs do not mask biological diversity under arbitrarily defined representative sequences (Selosse et al., 2016).

A detailed description of the taxonomic assignment and trophic guild assignment procedures can be found in the supplementary information document (Method S1). Briefly, taxonomic identity was assigned with the RDP naïve Bayesian classifier with a confidence threshold of 0.8 (Wang et al., 2007). We employed UNITE’s SH database from February 2020 as a reference (DOI: 10.15156/BIO/786368, Nilsson et al., 2019). Unidentified sequences were subjected to a second classification attempt with the blast+ algorithm (Camacho et al., 2009). This second query was restricted to version 5 of NCBI’s database of eukaryote ITS reference sequences (O’Leary et al., 2016). Guild and trophic mode of variants was retrieved from FUNGuild database (Nguyen et al., 2016). The database was accessed in October 2020, retaining only those matches with probable or highly probable assignment confidence. Given many fungal taxa exhibited multiple possible trophic modes, inherited annotations were edited to generalize guild classification. Finally, growth morphology was also annotated (i.e. unicellular, filamentous or dark septate endophyte).

### Statistical analysis

Analysis was conducted in R version 3.6.3 using packages adespatial (Dray et al., 2019), ALDEx2 (Fernandes et al., 2014), DHARMa (Hartig, 2020), dplyr (Wickham et al., 2018), emmeans (Lenth et al., 2021), ggplot2 (Wickham, 2016), ggpubr (Kassambara, 2018), glmmTMB (Brooks et al., 2017), lme4 (Bates et al., 2015), phyloseq (McMurdie & Holmes, 2013), rslurm (Marchand et al., 2021) and vegan (Oksanen et al., 2018). Some analyses that required extensive computation resources were performed at the high performance computer cluster of Freie Universität Berlin (Curta, DOI:10.17169/refubium-26754).

### Organization of the data, normalization and description of root associated communities

Given databases are strongly biased towards specimens from temperate ecosystems (Khomich et al., 2018), many variants recovered here remained unclassified at the finer levels of the taxonomic hierarchy (i.e. family, genus or species). Therefore, variants recovered were either analyzed together or grouped according to the phylum to which these were assigned. It was assumed that taxonomic classification at the kingdom and phylum levels are robust to updates in the taxonomy or improvements of reference databases.

Unequal sequencing depth among samples was normalized by random subsampling without replacement (n = 100) to a minimum depth of 26 278 reads per sample. This is the minimum library size obtained here. Although rarefaction is an imperfect normalization method (McMurdie & Holmes, 2014), it remains a valid and widely applied method when analyzing presence-absence data (Weiss et al., 2017).

Statistical inferences of community level attributes (i.e. richness and community composition) focused on well represented phyla. That is, Ascomycota, Basidiomycota and Mortierellomycota. These three phyla will be hereafter referred to as focal phyla. Glomeromycota was not included among focal phyla, since we have analyzed the response of Glomeromycota communities to the same experimental manipulations previously with more appropriate primer sets (Camenzind et al., 2016; Dueñas et al., 2020).

### Effects of fertilization on alpha diversity of root associated fungal communities

Alpha diversity was characterized as the total number of fungal ASVs per sample (*SF*), and as the total number of ASVs corresponding to the focal fungal phyla (*SP*). These richness estimates were then used as response variables to investigate how taxonomic diversity changes in response to nutrient manipulation. Given richness estimates obtained here were overdispersed (i.e. variance was larger than the mean), and the variance was not homogeneous across treatments, data was modelled with generalized linear mixed effect models (Bolker et al., 2009).

Fungal richness in sample *k* within block *i* and plot *j* (*SFijk*) was modelled as a function of fertilization with nitrogen (*Nijk*, categorical, two levels), phosphorus (*Pijk*, categorical, two levels) and their interaction. *Plotj* nested within *Blocki* were specified as a nested random term with a random intercept. It was assumed that variance follows a negative binomial distribution and the link function was the natural log. The full model structure and associated assumptions are presented according to the notation suggested in (Zuur & Ieno, 2016) in Eq. 1:

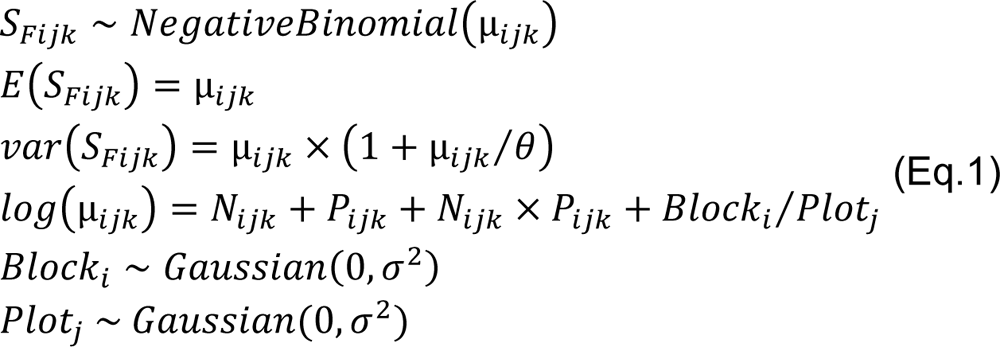

where *θ* is an overdispersion parameter allowing the variance to increase quadratically with the mean (Brooks et al., 2017). Richness at the phylum level (*SPijk*) was modelled maintaining essentially the same structure, except for the inclusion of a fixed term representing focal phyla (*Φijk*, categorical, three levels) and its interaction with fertilization factors. This term allowed to estimate whether different fertilization regimes affected the mean richness of focal phyla differently.

The magnitude and direction of fertilization effects on richness was determined by assessing the degree of overlap between the 95% confidence intervals (CI) of mean richness estimates between treatments and control (Hector, 2015; Nakagawa & Cuthill, 2007). Confidence intervals were computed by iteratively refitting models to parametric bootstraps of the original data (n=1000; Davison & Hinkley, 1997).

Fertilization effects on mean richness of focal phyla was assessed in the same way. In order to corroborate the inferences drawn from CI comparisons, *p* values were estimated from a Dunnett’s *t* test, thus correcting such values for multiple comparisons (Dunnett, 1955).

### Effect of fertilization on beta diversity of root associated fungal communities

All analyses in this section were conducted using normalized presence-absence ASV tables. ASV read counts were not taken into account for this analysis given such counts cannot be assumed to represent the underlying abundance of marker molecules in the environment (Gloor et al., 2017). In addition, given one sample failed (see Results section) and current implementations of restricted permutation designs only allow to conduct tests on balanced datasets, 15 samples were randomly excluded to achieve this balance (n=80).

Beta diversity was defined here as turnover of phylotypes across samples. To have an idea of how beta diversity of RAF communities in this site compares to fungal communities at other tropical forests, beta diversity was first summarized by the binary form of Jaccard dissimilarity index. This is an often employed dissimilarity index and was computed for both ASV and OTU tables. This was considered necessary to evaluate whether the estimates obtained employing ASVs are consistent to those obtained when employing OTUs. The strength and direction of the correlation between these two estimates was assessed through a Mantel correlation test (n=999). Similarly, to assess if distance between samples affected dissimilarity estimates, a Mantel correlation test was employed. Distance between samples was drawn from a Euclidean distance matrix of the UTM coordinates of each sample (coordinate reference system=WGS84, zone=17). The presence of a correlation between distance and dissimilarity would hint at the need to remove the trend imposed by distance (i.e. detrend) prior to testing for spatial autocorrelation.

Partial canonical redundancy analysis (RDA; Legendre & Legendre, 2012) was employed to model the variation in community composition of root associated communities as a function of fertilization. A total of four models were specified: one grouping all ASVs irrespective of their phylum assignation, and three additional ones corresponding to the focal fungal phyla examined in this study. ASV tables were specified as multivariate response variables while fertilization factors were specified as constraining variables. That is, a two-way factor model with interactions (i.e. ∼ *N*P*). To account for spatial dependencies, eigenvector based spatial analysis (Bauman et al., 2018; Dray et al., 2006) was employed. Spatial analysis allowed to identify positive autocorrelation signatures on the residuals of detrended ASV presence-absence tables (Borcard et al., 2018). When significant positive spatial autocorrelation was indeed detected, the correspondent Moran eigenvector maps (MEMs) were included in each model as partial spatial covariates. If no positive autocorrelation was found, only the spatial coordinates of each sample in UTMs were included as partial spatial covariates. Spatial autocorrelation is often used as an indicator of the presence of spatially clustered ecological processes that are structuring fungal communities.

To test if fertilization or spatial terms explain more variance in community composition than would be expected by chance, ANOVA like permutation tests (i.e. anova.cca()) were performed for each model (Legendre et al., 2011). Permutations were restricted within blocks to account for the experimental design structure. That is, free permutations (n = 9999) of data rows were allowed within blocks. Non metric multidimensional scaling (NMDS) was used to graphically represent dissimilarity between communities as a function of treatments. The NMDS algorithm was set to find an optimal four-dimensional solution employing 200 random starts. An additional run with default parameters starting from the solution arrived on the first set of iterations was used to plot the final ordinations.

Variance partitioning was employed to quantify the relative importance of covariate categories in structuring root associated fungal communities. Covariates were divided in three categories: fertilization (∼ *N*P*), spatial (*Long+Lat or MEMs*) and environmental. Environmental variables refer to plot wise measurements of organic layer pH, tree species richness, C:N ratio and resin available P. Details of the methods employed to measure these environmental variables can be found in Dueñas *et al*. (2020). A goodness-of-fit coefficient for each of these categories in every model was calculated. Given some categories encompassed more covariates than others, the adjusted coefficient of determination (adjusted R^2^) was calculated to correct for inflation of R^2^ (Peres-Neto et al., 2006).

### Sensitivity tests

Phylotype tables recovered by meta-barcoding are typically composed of a large number of features with low read counts, which are often suspected of being artifacts (Bálint et al., 2016). Thus to ascertain that the effects of experimental fertilization were not driven by its effects on rare ASVs only, all tests were repeated by segregating ASV tables by the frequency of occurrence of each feature. Frequent ASVs were defined as those present in at least 5 samples across the dataset. Consequently, rare ASVs were those observed in fewer than 5 samples.

### Differential abundance

Although read counts derived from high throughput sequencing cannot be assumed to represent the underlying abundance of a taxon in the environment, with proper statistical treatment, such information can be used to test if experimental manipulation systematically changes the read count of a given taxon (Gloor et al., 2017). Since conducting differential abundance testing at the level of ASVs is computationally prohibitive, it was investigated if the signal of an effect in read counts can be detected when grouping reads by their taxonomic assignment at phyla and orders, or by their trophic guild classification. In contrast to the analyses for community level attributes, all phyla were included in this analysis.

The package ALDEx2 (Fernandes et al., 2014) was employed. This package uses Bayesian methods to estimate test parameters while modeling technical variance. ALDEx2 first models the precision of count estimates via Monte Carlo (MC) resampling. Here, 1000 MC resampling iterations were specified. Values generated in each iteration were transformed to relative abundance by a center log ratio transformation (CLR). CLR is defined as the logarithm base 2 of the ratio between the abundance of feature *i* in sample *j* and the geometric mean of abundances in sample *j* (Fernandes et al., 2014; Gloor et al., 2017). Then the package can model if the variation in relative feature counts (*Cclr*) can be attributed to fertilization or to technical variation by means of generalized linear models (glms). Fertilization factors were specified as explanatory terms via a model matrix (*Cclr ∼ N*P*). A *t* tests of significance for each term in the model and a false discovery rate correction to the corresponding *p*-value (Benjamini-Hochberg) was estimated. Given a separate glm is fitted for each MC iteration, the mean of glm parameters across MC instances is reported.

## Results

### Sequencing results

A total of 95 samples were amplified and generated 17,338,558 reads. Sample 16F failed to amplify presumably because of low gDNA concentration (0.5 ng µl^-1^). After bioinformatic processing 6,712,425 remaining reads were mapped to 7,083 ASVs. Taxonomic classification routines confirmed 6,333 of these ASVs were likely of fungal origin. The remaining 700 ASVs predominantly matched plants and protists, thus were excluded from the dataset. Normalization by rarefaction discarded 539 variants, yet a rarefaction curve indicated that the estimated ASV richness per sample was close to saturation for most samples at the chosen normalization threshold (Fig. S2).

### Taxonomic and guild diversity

ASVs were placed within 10 phyla, 35 classes, 93 orders and 197 families. Classification efficiency declined with increased taxonomic resolution. While all fungal variants could be classified to phylum level, only 22.5% could be confidently identified to genus. In terms of read counts, Ascomycota and Basidiomycota were the most frequently recovered (66.3% and 30.5% respectively, Fig. 1a). By contrast, basal phyla together with phyla within subkingdom Mucoromyceta corresponded to only 3.3% of total reads. Beyond phylum level, Leotiomycetes and Sordariomycetes within Ascomycota accounted for the largest proportion of reads (Fig. 1b). By contrast, Agaricomycetes largely dominated the read proportion within Basidiomycota. Orders Helotiales, Sebacinales and Agaricales, concentrated an appreciable proportion of reads among orders (Fig. 1c). Archaeorhizomycetales was also present, albeit in low proportion.

**Figure 1.**
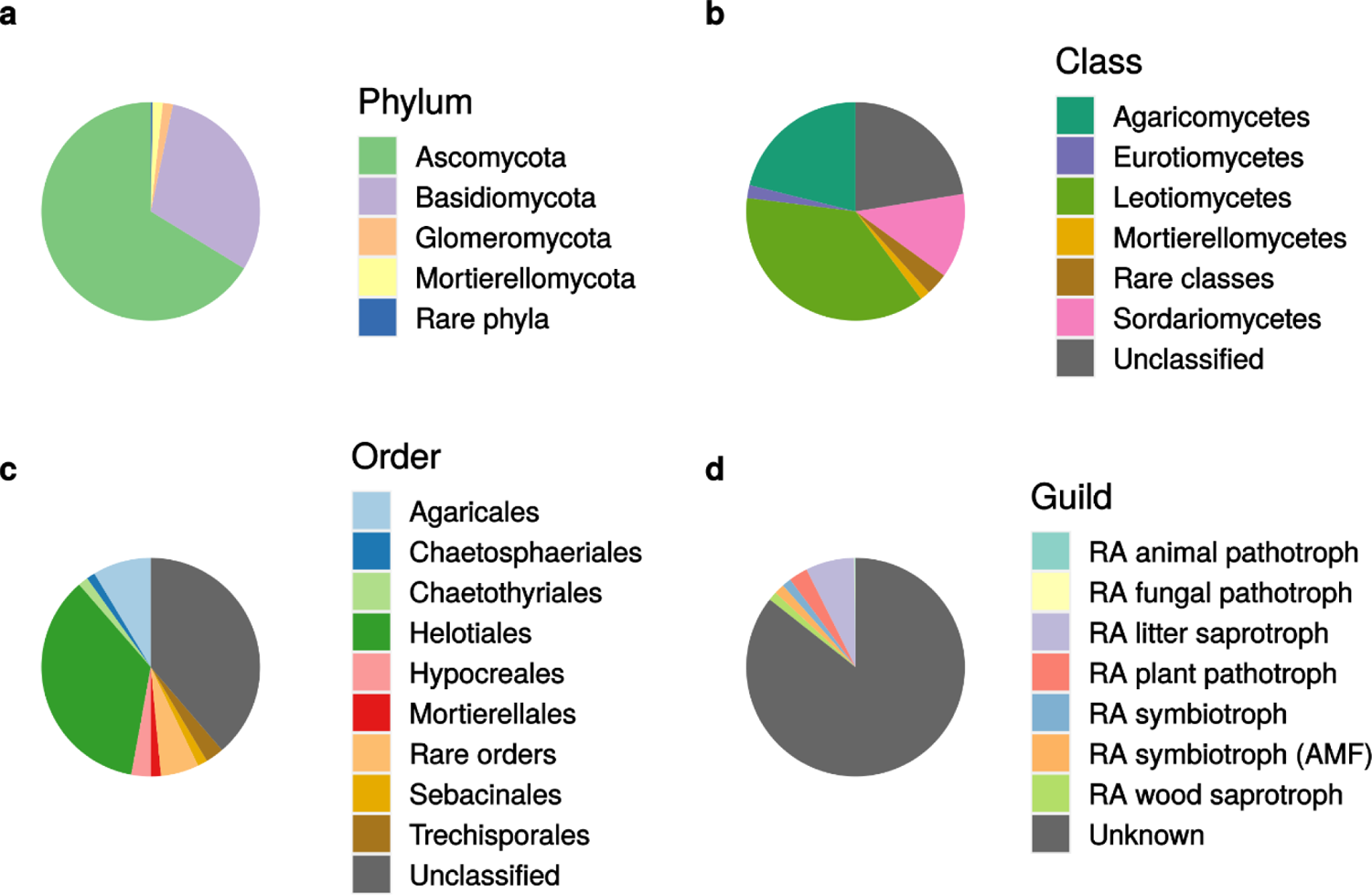
Relative abundance of amplicon sequence variants (ASVs) grouped according to their taxonomic classification at different levels (a,b,c), or according to their putative trophic guild (d). RA stands for root associated and AMF for arbuscular mycorrhizal fungi. Read depth was not normalized prior to plotting. Low read count clades, defined as those with proportional abundance of less than 1% of totals, were aggregated in one category across plots.

There was a wide variation in total ASV richness among phyla, with lineages within subkingdom Dikarya showing the highest diversity range (Basidiomycota– Ascomycota: 1729–4001 ASVs, Fig. 2b). Basal lineages in general exhibited low to very low taxonomic richness, with Mortierellomycota and Glomeromycota the most diverse among these (144 and 297 ASVs, respectively). By contrast, Mucoromycota, Rozellomycota, Zoophagomycota, Basidiobolomycota, Chytridiomycota and Kickxellomycota ASV richness was overall low (range: 3–77 ASVs).

**Figure 2.**
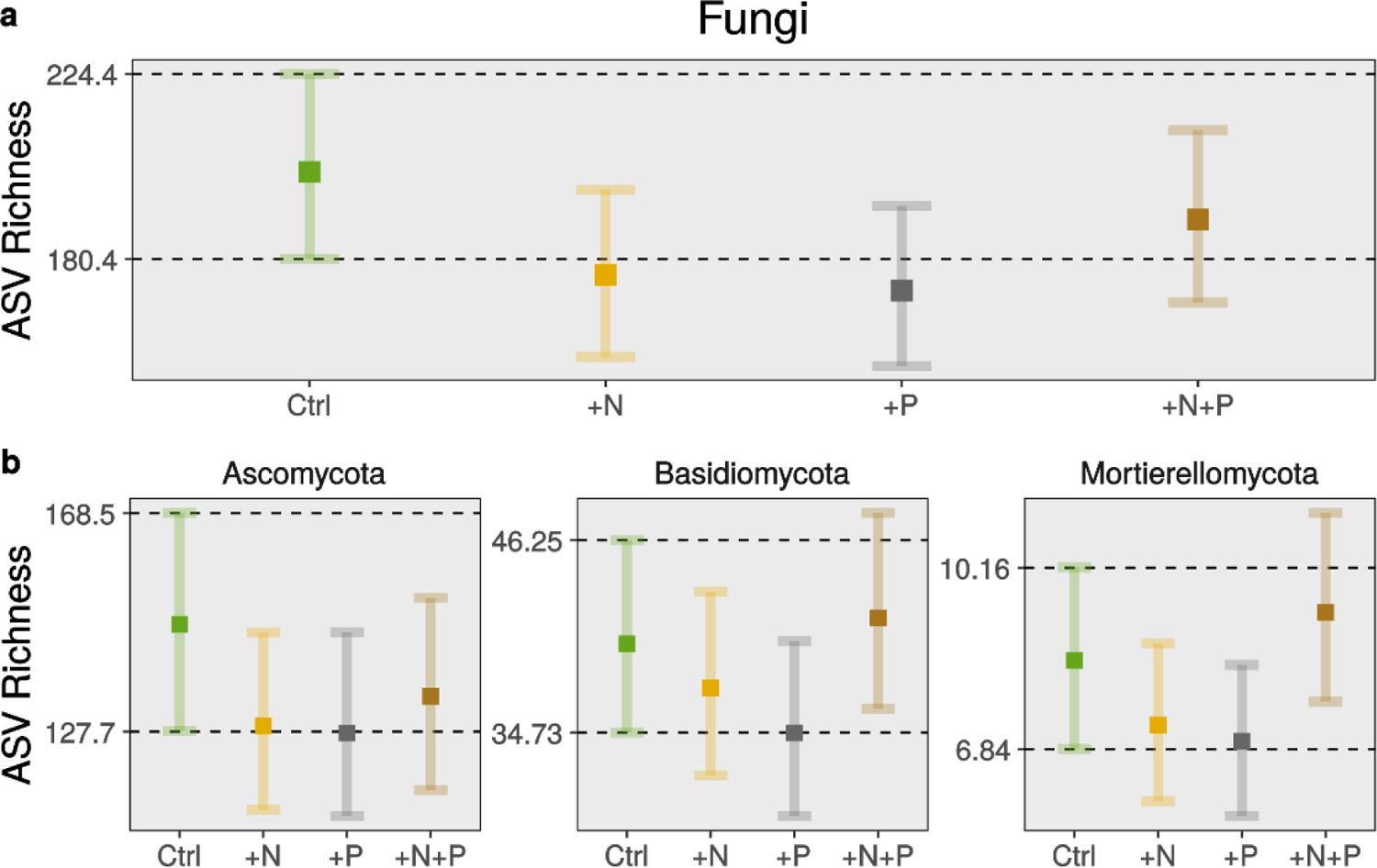
Effects of long term fertilization on taxonomic diversity of root associated fungal communities (n=95). Panel a depicts generalized linear mixed model (glmm) estimates of mean ASVs richness per sample (excluding Glomeromycota variants). Panel b shows estimates of mean ASV richness per sample of the most frequently recovered phyla. Solid squares represent model mean estimates. Whiskers represent 95% confidence intervals estimated by refitting glmms 1000 times with parametric bootstraps of the original data. Breaks and horizontal dashed lines in y-axis correspond to CIs estimates of controls. Prior to calculate ASV richness per sample, read depth was standardized across samples by rarefaction.

Classification into trophic guilds was sparse. Only 1155 variants (18.3%) inherited a guild classification. Litter saprobes was the guild with the largest proportion of read abundance (Fig. 1d), followed by plant pathotrophs and wood saprobes. Guild variant richness tracked this pattern, with litter saprobes being the richest (467 ASVs) followed by wood saprobes and plant pathotrophs (78–204 ASVs, respectively). Symbiotroph read abundance was comparable to that of pathotrophs; however, more than half of those reads mapped back to AMF variants (297 ASVs). Beyond AMF, considerably fewer reads and variants (39 ASVs) were annotated as putatively symbiotrophic. Among these, 30 variants corresponded to genera confirmed as ectomycorrhizal: *Cortinarius, Tomentella, Tellophora, Inocybe, Endogone* and *Clavulina*. Remarkably, a few variants annotated within genera with a suspected ectomycorrhizal status were also recovered (9 ASVs, *Serendipita* and *Hymenoscyphus*).

Regarding growth mode and morphology, a small proportion of reads (2.3%) corresponded to variants annotated as dark septate endophytes (70 ASVs). The most prevalent and abundant taxa among DSE were *Pezoloma ericae* and the genus *Oidiodendron*. By contrast, yeasts were less abundant in terms of read proportion (0.6% reads), yet more diverse than DSE (94 ASVs).

### Fertilization effects on alpha diversity

We observed that seven years of moderate fertilization rates did not elicit changes on RAF communities’ alpha diversity (Fig 2a, Table 1). Model estimates showed that, in relation to control, mean ASV richness decreased by 12.1% (mean, 95% CIs: 176.7 ASVs, 157.2–196.9) after N addition and by 14% (172.9 ASVs, 155.1–193.1) after P addition. Combined addition of N and P reduced mean ASV richness by 5.6% (189.8 ASVs, 170.1–211.1). A considerable overlap of CI estimates between fertilization treatments and control indicate that the probability such reductions could have been observed by random variation of the data is larger than 5% (Fig. 2a).

**Table 1.** Dunnett’s *t* test contrasting the difference in mean ASV richness between fertilization treatments and controls (n = 95). Dunnett procedure corrects *p* values to reduce false positives due to multiple contrast with the control. Only focal fungal clades recovered in this study were tested with the exception of Glomeromycota.

| Taxonomic level | Contrast | $t$ | $p$ |
| --- | --- | --- | --- |
| Fungi | + N vs. Ctrl. | -1.639 | 0.251 |
|  | + P vs. Ctrl. | -1.886 | 0.158 |
|  | + N + P vs. Ctrl. | -0.730 | 0.780 |
| Ascomycota | + N vs. Ctrl. | -1.404 | 0.364 |
|  | + P vs. Ctrl. | -1.500 | 0.313 |
|  | + N + P vs. Ctrl. | -0.975 | 0.629 |
| Basidiomycota | + N vs. Ctrl. | -0.648 | 0.825 |
|  | + P vs. Ctrl. | -1.338 | 0.401 |
|  | + N + P vs. Ctrl. | 0.361 | 0.946 |
| Mortierellomycota | + N vs. Ctrl. | -1.078 | 0.563 |
|  | + P vs. Ctrl. | -1.355 | 0.392 |
|  | + N + P vs. Ctrl. | 0.731 | 0.780 |

Dunnett’s *t* tests supported the CI assessment, confirming that no fertilization treatment elicited a statistically significant reduction in mean ASV richness at the kingdom level (Table 1).

Trends observed at the kingdom level remained consistent across focal phyla (Fig. 2b). That is, no fertilization regime changed mean ASV richness among these phyla. Dunnett’s *t* tests supported the CI assessments, showing none of the differences elicited by fertilization treatments were statistically significant (Table 1). Assumption validation plots and coefficients in the original scale for both models are presented in supplementary Figures S3–4 and Tables S1–2, respectively.

When the analysis was repeated segregating the data according to frequency of variants, trends remained similar for the kingdom level and for Ascomycota (Fig. S5a, b). Somewhat stronger mean richness reductions were observed amongst rare Basidiomycota and Mortierellomycota assemblages. However, Dunnett’s *t* tests indicated only mean richness of rare Mortierellomycota decreased significantly after N addition (*p* = 0.03) or P addition (*p* = 0.04), but not after the addition of both nutrients. In stark contrast to the overall trend, mean richness of rare Basidiomycota variants was higher than the mean richness of frequent ones (Fig. S5b).

### Fertilization effects on beta diversity

Beta diversity of RAF communities, as characterized by the Jaccard index, was large in this system (mean, ±SD: 0.92, ±0.03). This was the case irrespective of the resolution of taxonomic units employed (Fig. S6a). Dissimilarity of communities in terms of ASVs was strongly and positively correlated to the dissimilarity of communities constituted by OTUs (Mantel *ρ* = 0.83, *p* < 0.001). Overall, dissimilarity between communities constituted by ASVs was not correlated with distance (Mantel *ρ* = 0.13, *p* = 0.395). However, this pattern was inconsistent across treatments (Fig. S6b). While the correlation was weakly positive and significant among assemblages within Control, and +P plots, it was not significant when considering assemblages within the N addition treatments (Table S3).

Removal of samples to achieve a balanced design left a total of 5655 ASVs and 5 samples per experimental plot (n = 80). Spatial eigenvector analysis showed that the residuals of presence-absence tables were spatially independent in most cases (Table S4). Positive spatial autocorrelation among residuals was found only for assemblages consisting of frequent fungal variants and for assemblages consisting of all or frequent Ascomycota variants. Consequently, MEMs as spatial covariates were included only for these three cases. In all other instances, UTM coordinates were specified as spatial covariates.

Permutation tests showed that both the addition of N and P explain a statistically significant portion of the variation in composition of root associated fungal communities (Table 2, Fig. 3). Test at the phylum level were rather inconsistent, with the exception of Ascomycota assemblages, for which the addition of N or P elicited significant community structure shifts. By contrast, assemblages consisting of Basidiomycota or Mortierellomycota variants responded weakly or not at all to fertilization with N or P. Specifically, composition of Basidiomycota communities shifted in relation to control only when P was added, while Mortierellomycota communities did not shift after any fertilization intervention. These results were robust to the inclusion of spatial covariates or when segregating data by variant occurrence frequency (Table S5). Yet two notable differences arose when analyzing only rare ASVs. The addition of N did not shift the structure of any of the clades considered and, the addition of P did affect the structure of the two most frequent clades recovered in this study (i.e. Ascomycota and Basidiomycota).

**Table 2.** Pseudo-*F* tests assessing whether the proportion of variance in community composition explained by fertilization factors is statistically significant. Spatial covariates were partialled out in all RDA models. Only the best represented fungal clades recovered in this study were tested with the exception of Glomeromycota. Values of *p* were calculated with permutation tests (n = 9999). Free permutations

| Taxonomic level | Factor | Pseudo *F* | *p* |
| --- | --- | --- | --- |
| Fungi | +N | 1.136 | 0.020 |
|  | +P | 1.441 | 0.000 |
|  | +N+P | 1.096 | 0.043 |
| Ascomycota | +N | 1.300 | 0.000 |
|  | +P | 1.404 | 0.000 |
|  | +N+P | 1.056 | 0.167 |
| Basidiomycota | +N | 1.046 | 0.224 |
|  | +P | 1.173 | 0.007 |
|  | +N+P | 1.114 | 0.053 |
| Mortierellomycota | +N | 1.164 | 0.164 |
|  | +P | 1.381 | 0.071 |
|  | +N+P | 1.328 | 0.099 |

**Figure 3.**
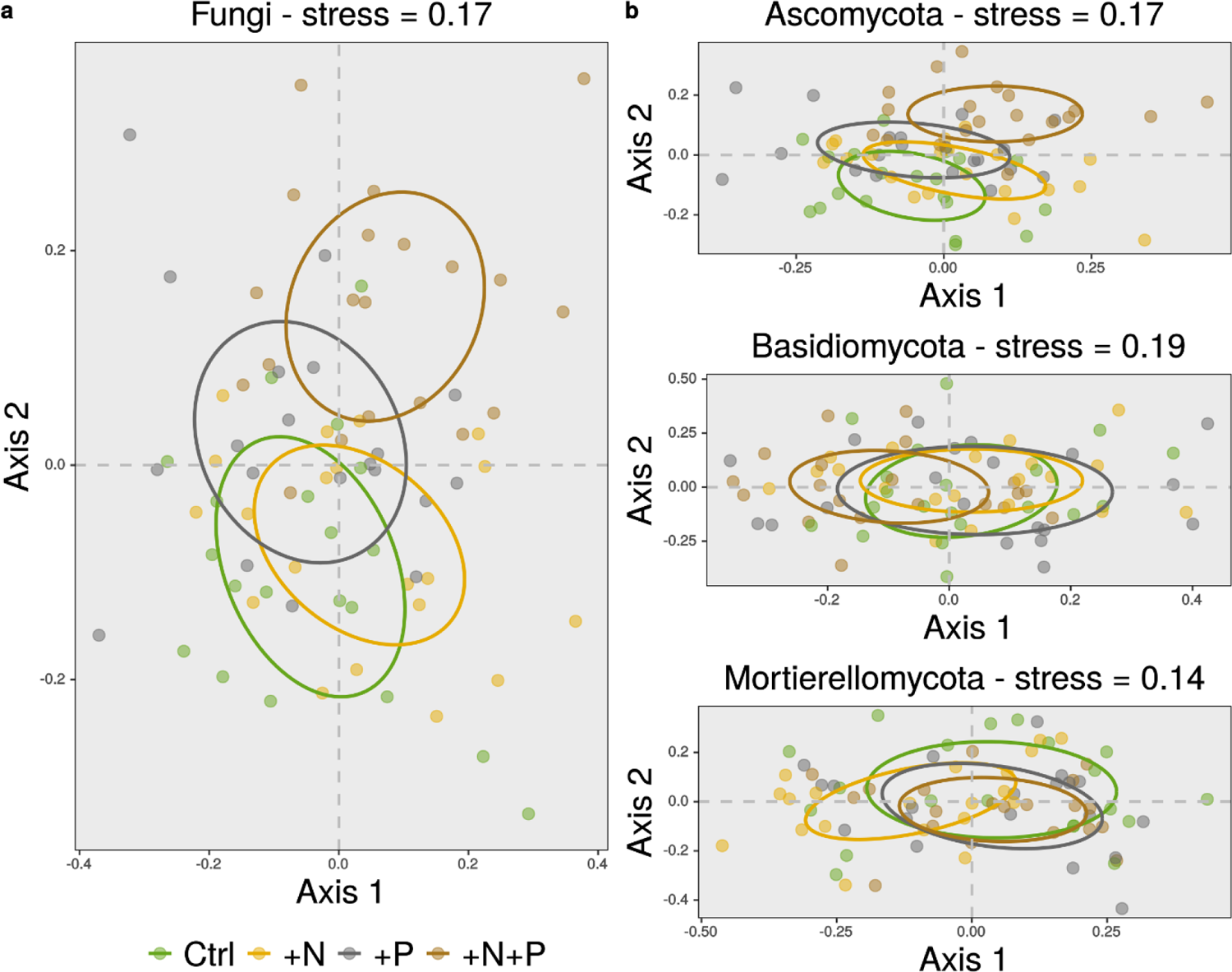
Non-metric multidimensional scaling (NMDS) of binary Jaccard dissimilarities (n=80). The NMDS algorithm was set to resolve ordinations with 200 random starts. Four dimensions were necessary to arrive to a fair representation of the dissimilarity matrix in the ordination space. However, only the first two dimensions are illustrated here. Each colored point represents a fungal assemblage. Ellipses represent one standard deviation from the group centroid.

Variance partitioning showed that none of the covariate groups analyzed contributed to explain a considerable portion of RAF community composition variability (Fig. 4). Fertilization factors had relatively low explanatory power, explaining the largest portion of variability among Ascomycota assemblages (range: 0.3–0.8%). Similarly, spatial covariates were generally poor predictors of RAF community structure, again with the exception of Ascomycota assemblages (range: 0.3–1.2%). Plot wise environmental covariates, such as pH, plant available P, C:N ratio and tree species richness boosted the explanatory power of multivariate models in some cases. For instance, environmental covariates appear to be better predictors of Mortierellomycota assemblage structure than either fertilization or spatial covariates (1.6% versus 0.4% and 0.7% respectively). The patterns just described varied widely according to whether ASVs were analyzed together or grouped according to phyla or frequency of occurrence. Regarding focal phyla, covariate groups cumulatively explained 1.4% of the variability in structure of Basidiomycota assemblages while these same predictors explained more than double of this proportion among Mortierellomycota assemblages (4.2%). Regarding occurrence frequency, covariate groups clearly explained a greater proportion of variance when assemblages where composed of frequent ASVs only. Conversely, predictor groups collectively explained less than 1% variability when considering assemblages exclusively composed of rare variants. The partitioning function found a large proportion of shared explained variability amongst the covariate groups analyzed.

**Figure 4.**
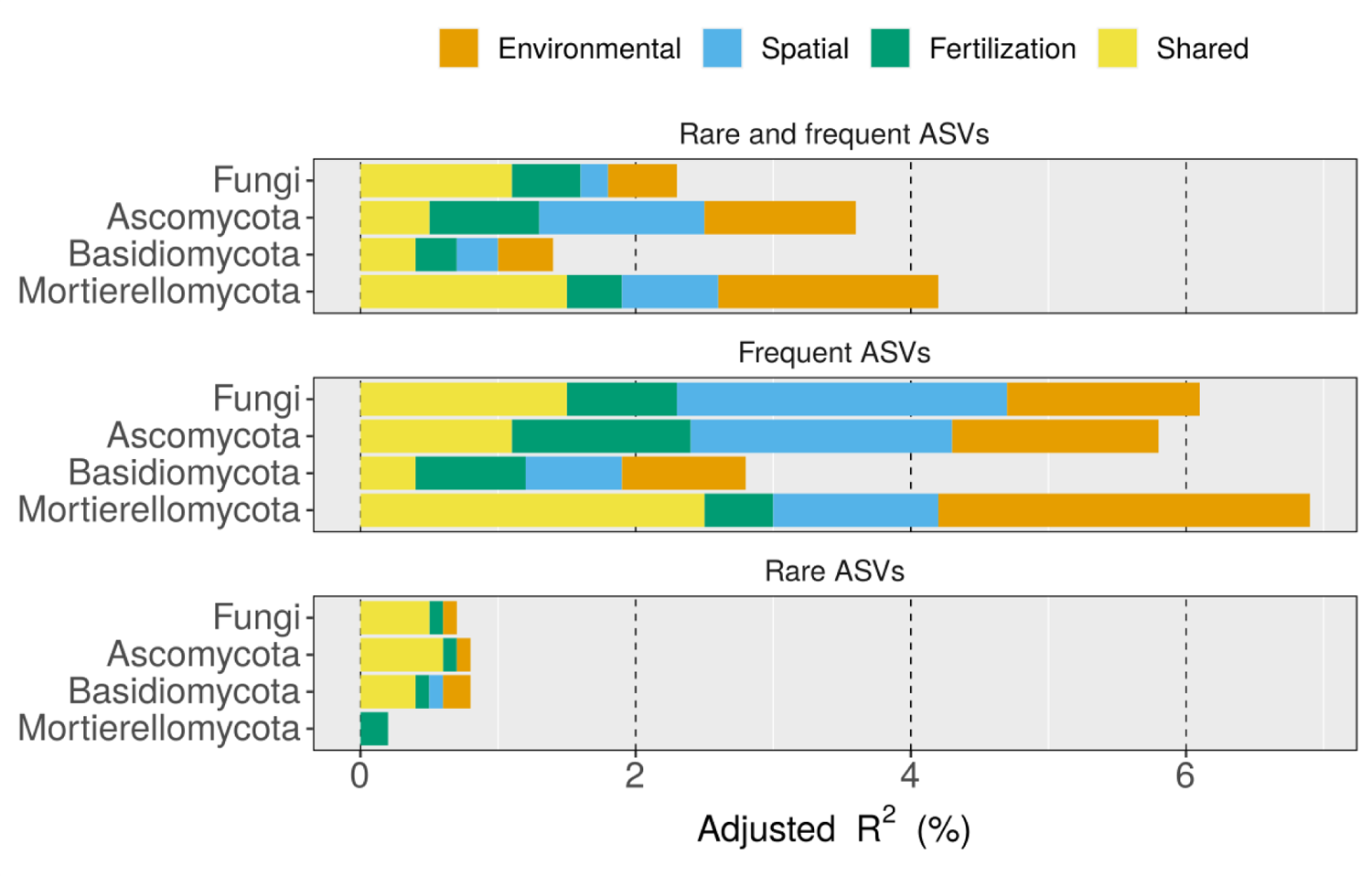
Proportion of variability in community composition explained by fertilization treatments among the focal fungal clades recovered in this study. The adjusted coefficient of determination (Adjusted R^2^) was estimated with a variance partitioning function and then transformed to a percentage for plotting. ‘Fertilization’ represents the variance explained by fertilization covariates. ‘Space’ represents the variation explained either by spatial coordinates or Moran eigenvector maps. ‘Environmental’ represent the variance explained by additional environmental variables. That is, plot wise soil pH, plant available P, C:N ratio and tree species richness. The magnitude of variance explained by each covariate group was estimated excluding all other covariate groups.

### Differential abundance

Relative abundance of reads amongst the great majority of phyla, orders or guilds did not change in response to fertilization (Fig. 5). Univariate tests indicated technical variation was as great as variation between treatments in almost all cases (Fig. S7). Few departures from this pattern were observed. For instance, the relative abundance of the order Glomerales was reduced while that of Sebacinales increased after the addition of P, yet univariate tests indicated such changes were marginally or non-significant after applying the B-H false positive correction (p = 0.08 and 0.18, respectively, Fig. S7).

**Figure 5.**
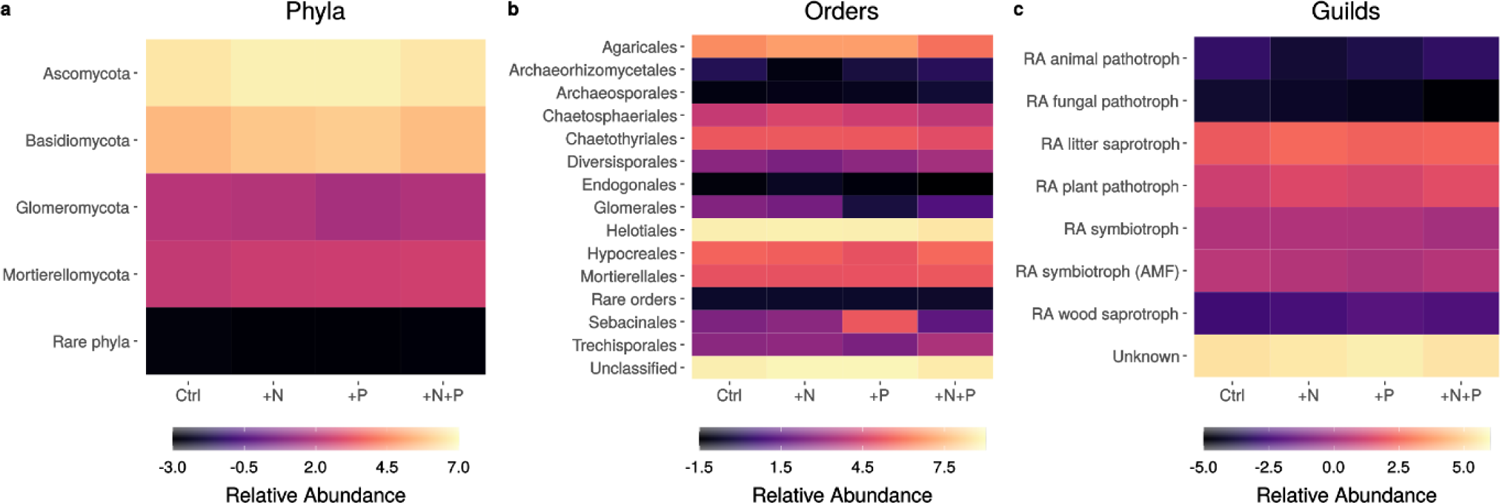
Heatmaps representing the change in mean relative read abundance of features as a function of fertilization treatments. Features refer to taxonomic clades and fungal guilds to which reads were mapped to. Prior to plotting, read abundance was transformed by the center log2 ratio transformation (CLR) to correct distortions in read counts. Lighter colors represent higher mean relative abundances, while darker colors represent lower relative abundances.

## Discussion

We characterized, for the first time via a deep sequencing approach, the non-mycorrhizal fungi associated with the fine root fraction of an Andean tropical forest. Our assessment revealed that the fine roots of tropical montane trees are not only associated with AMF variants, but with a diverse set of fungal variants, among which many are primarily known as saprobes. We have also shown that the attributes of RAF communities remained mostly stable after a simulated chronic increase in the input of P and N to the soil. That is, mean richness reductions following fertilization were neither generalized among fungal lineages nor clearly distinguishable to what would be expected by chance in most cases. Regarding community structure, though we observed an overall shift following N and P additions, at the phylum level these shifts were most prominent among Ascomycota assemblages, but inconsistently observed among other phyla. Furthermore, the variability explained by fertilization factors was overall low. Given the inconsistent and rather weak responses observed, we suggest that tropical montane RAF communities are robust to moderate increases in the atmospheric inputs of N and P. RAF communities in this system are dominated by phylotypes from sub-kingdom Dikarya. The overwhelming dominance of Ascomycota and Basidiomycota was expected, given it has been repeatedly reported for various neo-tropical lowland (Barberán et al., 2015; Peay et al., 2013; Schappe et al., 2017), and montane forests sites (Geml et al., 2014; Looby et al., 2016; Nottingham, Fierer, et al., 2018). While at the stand level similar dominance patterns have been reported for RAF communities (Schroeder et al., 2019), tree species level reports of communities dominated by Ascomycota (*Alnus accuminata*, Wicaksono et al., 2017) or by Basidiomycota (*Oreomunnea mexicana*, Corrales et al., 2017) also exist. Beyond the phylum level, our observation that Leotiomycete and Sordariomycete were abundant is consistent with a recent characterization of species within these classes as frequent components of soil fungal communities in mesic tropical forests (Egidi et al., 2019). As far as we know, Sebacinales associating with trees from the region have never been reported. Yet their presence is unsurprising given their cosmopolitan distribution, broad host-spectrum and trophic flexibility (Riess et al., 2014; Weiß et al., 2016). We believe Sebacinales presence, as well as that of Archaeorrhizomycetales deserve independent confirmation in order to establish if these play a beneficial role in the mineral nutrition of tropical montane trees (Pinto-Figueroa et al., 2019; Weiß et al., 2016).

A comprehensive characterization of trophic guild structure of RAF communities was not possible due to the large proportion of undescribed taxa and the lack of trophic guild information for tropical taxa. Yet we complemented the observations of early meta-barcoding studies of the fine root fraction of sub-tropical (Toju et al., 2014), and neo-tropical trees (Schroeder et al., 2019), which have reported the co-occurrence of symbiotrophs with a complex array of root associated guilds. The observation that the most frequent guilds in this fraction were saprobes, supports the idea that many members of RAF communities are horizontally transmitted and facultative (Hardoim et al., 2015; Rodriguez et al., 2009). An often given explanation to this pattern is that saprobic fungi can asymptomatically transition to an endophytic lifestyle, in order to ‘escape’ limiting conditions in other substrates (Baldrian, 2017; Nelson et al., 2020). Regarding non-AMF symbiotrophs, the recovery of several ecto- and ericoid mycorrhizal lineages in our communities is in line with the idea that soil and climatic variables drive their distribution, rather than host availability (Corrales et al., 2018; Peay et al., 2015; Wicaksono et al., 2017). However, it is worth noting that tree species suspect of broader mycorrhizal associations do occur in our experimental plots (e.g. *Bejaria aestuans* or *Graffenrieda emarginata*). Our study thus supports early assessments which reported morphological and molecular evidence of the presence of non-AMF mycorrhizal fungi in Andean forests (Haug et al., 2004; Kottke et al., 2004). The possibility that pathotrophs are as diverse and abundant as symbiotrophs in the fine root fraction is intriguing. Theory predicts that pathogen-mediated negative feedback balances the positive effects of mutualism thereby promoting the enormous plant biodiversity observed in the tropics (Bachelot et al., 2015). Perhaps unlike seedlings (Bachelot et al., 2017), adult trees have managed to escape negative feedbacks by establishing associations with a greater diversity of both mycorrhizal and non-mycorrhizal symbiotrophs. While these patterns must be interpreted with caution given the trophic guild of a fungal root associate is hard to predict from the identity of the taxa alone (Griffin & Carson, 2018), our attempt at characterizing fungal guild structure in this system has revealed a number of interesting patterns that await further confirmation.

RAF communities recovered in this study exhibited moderate richness levels. Providing context to this observation is challenging given the methodological inconsistencies of meta-barcoding studies (Bálint et al., 2016). However, a rough comparison of mean richness estimates (i.e. using the same number of samples, and clustering ASV into OTUs at 97% similarity) obtained at various neo-tropical forests suggests RAF communities in tropical Andean forests are less diverse than fungal communities found in soil or at lowland tropical forests (Table 3). This is in line with the metabolic theory of ecology which posits that richness decreases monotonically with temperature (Zhou et al., 2016), and fits well with the patterns revealed by the few fungal surveys conducted along elevation transects in wet tropical montane forests (Looby et al., 2016; Nottingham, Fierer, et al., 2018). This comparison also suggest that RAF communities are populated by a subset of the taxa found in soil, which fits the patterns revealed by recent surveys in temperate forests (Goldmann et al., 2016). We acknowledge, however, that until the richness of soil fungal communities at the same site, and at lower elevations, are thoroughly characterized, the question of how diversity estimates found here compare to other soil compartments and elevation belts will remain open (Looby & Martin, 2020).

**Table 3.** Estimates of mean fungal richness (*Sf*) and standard deviations (*σ*) across different types of wet neo-tropical forests. Estimates were obtained by randomly selecting nine samples from each site, as this was the minimum sample size of the studies compared. When appropriate, the random draw was selected from control plots only. All studies employed Illumina MiSeq platform and targeted the ITS region. For the purpose of this comparison, phylotypes were clustered as OTUs at 97% similarity and singletons (i.e. phylotypes occurring in only one sample) were eliminated.

| Sample type | Site | Forest elevation | $S_f$ | $\sigma$ | Reference |
| --- | --- | --- | --- | --- | --- |
| Composite soil | Panama (BCI) | Lowland | 183.44 | 58.09 | Barberán et al. 2015 |
|  | Panama (Campo Chagres) | Lowland | 449.67 | 68.85 | Schappe et al. 2017 |
|  | Panama (Pipeline) | Lowland | 434.67 | 67.95 | Schappe et al. 2017 |
|  | Panama (Santa Rita) | Lowland | 413.89 | 84.77 | Schappe et al. 2017 |
|  | Costa Rica (Monteverde) | Montane | 473.78 | 83.99 | Looby et al. 2016 |
| Mixed root | Mexico (Los Tuxlas) | Lowland | 497.33 | 107.66 | Schroeder et al. 2019 |
|  | **Ecuador (San Francisco)** | **Montane** | **107.78** | **14.29** | **This study** |

Contrary to our expectations, increased nutrient availability did not alter the mean richness of RAF communities, hence we did not find support for the resource ratio model expectations (Cardinale et al., 2009). Fungal richness has been shown to increase after long-term N addition in temperate forests soils (Morrison et al., 2016), and after long-term N and P additions to the organic layer of tropical forests (Kerekes et al., 2013). While it would be tempting to attribute such shifts only to the high fertilization rates applied (≥ 100 kg N ha^-1^ yr^-1^), comprehensive multi-site nutrient manipulation experiments (Leff et al., 2015) and meta-analysis (Zhou et al., 2020) indicate that, while there is considerable local variation, diversity indices of non-mycorrhizal fungal communities change little after fertilization treatments. Such neutral responses contrast with the clear richness reductions often observed among tropical forests AMF communities, even after moderate rates of N and P addition (Camenzind et al., 2014; Dueñas et al., 2020; Sheldrake et al., 2018). Yet they are consistent with the neutral richness responses observed among tropical ectomycorrhizal fungal communities (Corrales et al., 2017) after high N fertilization rates. Beyond the fertilization rate applied, what this evidence seems to indicate is that diversity of different fungal functional groups exhibit distinct sensitivities to soil eutrophication. Such sensitivity appears to depend on the ambient physico-chemical conditions of the soil. For instance, in the fertilization experiment surveyed by Kerekes *et al*. (2013), the largest richness gains were observed in the combined P and K addition treatment, which constitute two of the most limiting resources in lowland tropical forest soils. Similarly, Morrison *et al*. (2016) reported significant richness gains after N additions, in a system where N is the most limiting nutrient. Hence it appears that because Andean tropical forests soils have different nutrient limitations than temperate or lowland tropical systems, moderate N and P additions did not result in large richness increases or decreases.

We also hypothesized that nutrient imbalances caused by nutrient addition would elicit RAF community composition shifts. While we found some support for this prediction, community composition of most focal fungal phyla recovered here was not altered by the fertilization treatments. Moreover, our approach to measure changes in relative read abundance among focal phyla, orders and trophic guilds revealed that these remained stable across treatments. These results are in line with the notion that global change factors, and in particular nutrient deposition, are not always detrimental to microbial communities (Zhou et al., 2020). Indeed, the structure of fungal communities has been shown to respond in a variety of ways to increments in nutrient availability. While many studies in temperate and boreal forests report shifts in community composition, as well as changes in the abundance of certain lineages as a result of N addition (Edwards & Zak, 2011; Maaroufi et al., 2019; Morrison et al., 2016), reports of neutral responses in these indicators are not uncommon (Carrara et al., 2018; Zak et al., 2019). A similarly inconsistent response appears among the few studies conducted at tropical forests. Reports range from shifts in community composition and increases in PLFA markers as a result of N addition (Corrales et al., 2017; Cusack et al., 2011; Nottingham, Hicks, et al., 2018), to mild shifts in community structure or PLFA abundance after the experimental addition of N or P (Kaspari et al., 2010; Kerekes et al., 2013; Krashevska et al., 2014). Our results thus add to this variability and highlight the fact that more tropical sites need to be assessed in the context of soil eutrophication in order for robust patters to emerge.

Many features of fungal communities can allow them to resist the effects of moderate increases in nutrient availability. Since deep sequencing approaches cannot distinguish between the active and the dormant fractions of fungal communities (Fierer, 2017), it is possible that many members of the community are able to persist in a dormant status (Blagodatskaya & Kuzyakov, 2013). Such portion of the community will still require to consume maintenance resources, yet contributes directly to the resistance of the system because it allows many taxa to persist despite an unfavorable shift in environmental conditions (Joergensen & Wichern, 2018). In addition to opt for dormancy, the active portion of fungal communities can adjust their enzymatic profile in response to long term nutrient additions without necessarily reflecting such functional shifts as major changes in richness or community composition (Edwards & Zak, 2011; Morrison et al., 2018; Zak et al., 2019). This is an important point, because it implies that a lack of change in fungal community attributes does not necessarily translate in functional stability. In addition to these possibilities, recent experimental evidence has shown that soil fungi exhibit extremely flexible stoichiometric homeostasis in relation to the stoichiometric ratio of culturing media (Camenzind et al., 2021). If this ability is confirmed and widespread across the fungal kingdom, it will mean that non-mycorrhizal fungi have yet another possibility to bridge resource gaps in time or space, thus allowing taxa to persist despite shifts in the nutrient supply. In summary, given fungi possess a unique variety of nutrient acquisition strategies and life history traits (Treseder & Lennon, 2015), a diverse fungal community is robust to moderate increases in nutrient availability given its constituents have the potential to compensate functionally or to become asynchronously active (Morrison et al., 2018).

It is also possible that RAF communities are simply not structured by the same drivers affecting soil fungal or mycorrhizal communities. We observed high beta diversity, which was weakly correlated with distance between samples. This is remarkable, given the small spatial extent covered in this study (0.64 ha). Indeed, high turnover rates at local scales appear to be a generic feature of soil fungal communities (Kaspari et al., 2010; Zinger et al., 2019), as well as of leaf associated fungal communities (Meiser et al., 2014) in neo-tropical forests. This suggests that drivers other than soil conditions are structuring fungal communities and in particular plant associated communities. For instance, RAF communities could be structured by the chemical and physical characteristics of the fine root fraction (Nguyen et al., 2020). It is also reasonable to expect plant-fungal interactions to leave a signature in RAF community structure (Schroeder et al., 2019). Both interpretations are in line with the strong correlations between tree and fungal community composition reported at some lowland tropical forests (Barberán et al., 2015; Peay et al., 2013). Although the root environment characteristics and host filters appear to be relevant for RAF communities, we cannot exclude that small scale variation in soil physico-chemical properties or even stochastic factors (i.e. ecological drift *sensu* Vellend, 2016) are playing a role in structuring these communities. Our results, however, seem to balance against the influence of soil environmental filters, given positive spatial autocorrelation was inconsistently observed amongst the communities studied here (Table S4). It is thus clear that more research on the multiple assembly rules that could drive RAF community structure is needed before we can arrive at solid conclusions about the relative importance of each one of these factors.

In conclusion, we found no evidence that indicates a seven year long moderate increase in nitrogen and phosphorus supply affects RAF community richness in this Andean forest. RAF community composition on the other hand is moderately affected by these treatments, driven mostly by changes observed among Ascomycota assemblages. RAF within Ascomycota thus appear to be the most responsive clade to soil eutrophication. Altogether this indicates that the constituents of RAF communities, particularly those belonging to recently evolved, non-mycorrhizal fungal lineages, are quite adaptable to changes in the soil nutrient supply. This flexibility in turn is expressed as robustness of community level attributes to nutrient addition. As Kaspari *et al*. (2010) stated in a rather clever adaptation of Beijerinck’s conjecture, it appears that in highly diverse tropical forests, fungi are “never everywhere—but plastic and adaptable”. Our data support this idea and also suggests that the root environmental characteristics and plant top down control contribute to shape the structure of RAF communities. While these results are encouraging, we still need to tackle the question of how small scale gradients in soil and root nutrient availability structure RAF communities. Moreover, we need a better understanding of how biotic interactions shape these communities and how all this complexity affects the ecosystem level response to global change drivers (Carrara et al., 2018). Yet our most pressing priority is to continue to close the wide gap in our understanding of tropical fungal communities in relation to temperate or boreal ecosystems. The global implications of climate change and the disproportionate contribution of tropical biomes to dampen or enhance these effects make this a crucial task.

## Acknowledgements

We acknowledge DFG (grant no. PAK 823/1) for financial support and thank the Ecuadorian Ministry of the Environment for granting the necessary permits to conduct this study. We extend our gratitude to Milton Ortega and the staff of Universidad Técnica Particular de Loja and Naturaleza y Cultura foundation for assistance during field campaigns. We thank Macrogen Inc. for their assistance with sequencing. Finally, we acknowledge Catlin Looby for making available data from prior publications upon request.

## Author contributions

JFD performed DNA extractions, conducted bioinformatics and statistical analysis and wrote the first full draft of the manuscript. TC collected field samples and assisted with statistical analysis and manuscript write up. SH assisted with sequencing work. JH designed the study, conducted soil chemical analysis and taxonomic identification of trees. JPS assisted with field collection. MCR designed the study, obtained financial support and contributed with ideas for the analysis. All authors revised, edited and approved the final version of the manuscript. The authors declare no conflict of interest.

## Data availability

All sequences derived from this study are available at the European Nucleotide Archive (ENA). ASV sequences can be found with accessions OU000611– OU006943. Raw reads are available with accessions ERX5565434–ERX5565528. Sample metadata is available with accessions ERS6425851–ERS6425946.

