## Supplementary Information for "Non-mycorrhizal root associated fungi of a tropical montane forest are relatively robust to the long-term addition of moderate rates of nitrogen and phosphorus"

**Contents:**

**Method S1** Details of the taxonomic assignation and trophic guild assignation procedures.

**Figure S1** Map of the nutrient manipulation experiment and sampling design.

**Figure S2**. ASV accumulation curve.

**Table S1** Coefficients of the first generalized linear mixed effect model (glmm).

**Figure S3** Assumption validation of first glmm with the DHARMa method.

**Table S2** Coefficients of the second glmm.

**Figure S4** Second glmm assumption validation with the DHARMa method.

**Figure S5** Comparison of observed differences in ASV richness as a function of fertilization between frequent and rare variants.

**Figure S6** Beta diversity of root associated fungal communities.

**Table S3** Mantel correlation tests.

**Table S4** Tests of different neighborhood matrices to draw spatial eigenvectors.

**Table S5** Sensitivity tests of the effects of long term fertilization on the structure of root associated fungal communities.

**Figure S7** Graphical illustration of the results of univariate tests of the effect of fertilization treatments on the relative read abundance.

Non peer reviewed

**Method S1** Details of the taxonomic assignation and trophic guild assignation procedures.

Taxonomic identity was assigned with the RDP naïve Bayesian classifier with a confidence threshold of 0.8 (Wang et al., 2007). Such a high confidence score was selected to avoid forcing the classification of variants potentially representing novel or unknown organisms (i.e. over-classification). We employed UNITE's SH database from February 2020 as a reference (DOI: 10.15156/BIO/786368, Nilsson et al., 2019). As UNITE databases employ the taxonomic framework proposed by Tedersoo et al. (2018), this framework was also adopted here. This taxonomic framework is mostly consistent with the one in use by the International nucleotide sequence database (INSD) but proposes a series of updates to the higher level taxonomic nomenclature of the kingdom fungi. The most relevant changes to the present study are the elevation of Basidiobolomycota, Glomeromycota, Kickxellomycota and Mortierellomycota to the status of phylum; as well as changes in several classes within phyla.

Unidentified sequences were subjected to a second classification attempt. The blast+ algorithm was employed (Camacho et al., 2009), restricting the query to version 5 of NCBI's database of eukaryote ITS reference sequences (O'Leary et al., 2016). Only the 10 best matches with an e-value < 1e-10 were recorded. Matches were sorted in decreasing order by the percentages of query coverage and identity. Ultimately, only the highest ranked match per query sequence was preserved as the best. If query cover of the best match was less or equal to 90%, only domain and phylum names were annotated. If query cover was greater than 90% and identity percentage was greater or equal to 97%, taxonomic names were inherited up to species level. If identity was >= 95%, taxonomic names were inherited up to genus

Non peer reviewed

level. From there on up to family when identity was >= 90%, to order when identity was >= 80% and to class when identity >= 70%.

Guild and trophic mode of variants was retrieved from FUNGuild database (Nguyen et al., 2016). The database was accessed in October 2020, retaining only those matches with probable or highly probable assignation confidence. Given many fungal taxa exhibited multiple possible trophic modes, inherited annotations were edited to generalize guild classification. Thus, taxa that were not exclusively classified as mycorrhizal, fell into the general category of root associated (RA). Within this general category, several alternative trophic modes were emphasized. Finally, growth morphology was also annotated (i.e. unicellular, filamentous or dark septate endophyte).

Non peer reviewed

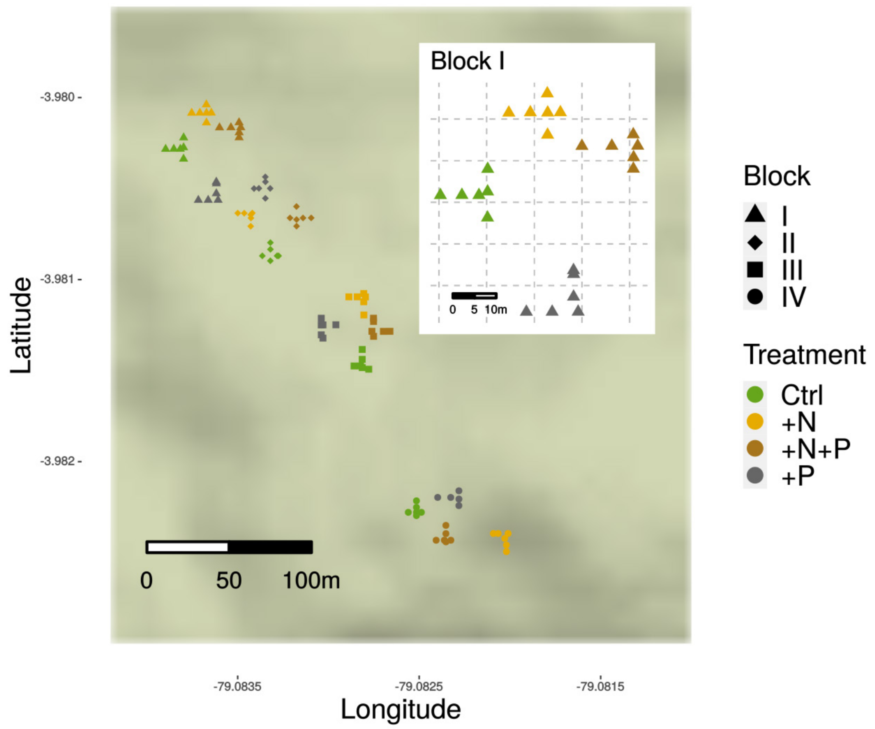

Figure S1 Map of the Ecuadorian nutrient manipulation experiment (NUMEX). Each colored symbol represents a mixed root sample within an experimental plot. Colors represent the different fertilization regimes to which plots were subjected to. Symbols indicate the spatial position of the four blocks. The inset details how plots are arranged within blocks, and the spatial distribution of samples along the two orthogonal transects within each plot. The experimental area amounts to 6400 m2. The shortest distance between samples was 1.01 m while the longest was 328.80 m. The terrain polygon layer was obtained from Open Street Maps.

Non peer

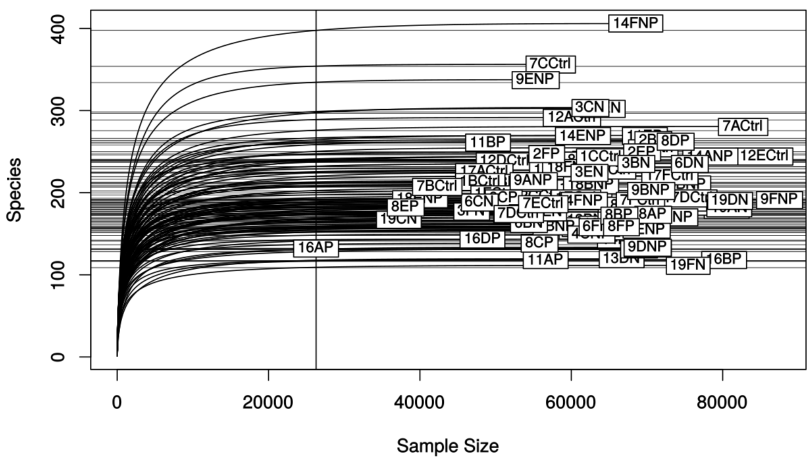

**Figure S2** ASV richness accumulation curve as a function of the number of reads per sample. The vertical line signals the minimum read count achieved in our dataset (26 278 reads). This is the cutoff value employed subsequently for normalization of sample read depth. Horizontal lines indicate the rarefied ASV richness estimated for each sample at the chosen normalization depth. Normalization was carried out by random sampling without replacement (n = 100). Unless otherwise stated, all results presented in the main text or below were estimated using the normalized dataset.

Non peer reviewed

**Table S1** Coefficients of the first generalized linear mixed effect model where fungal ASV richness was modelled as a function of fertilization treatments. The intercept estimate corresponds to the mean ASV richness for control plots. Estimates and standard errors are presented in the natural logarithmic scale, as this was the specified link function for the negative binomial probability distribution. *Z* represents the Wald-*Z* statistic, while *p* represents the probability of observing a theoretical *Z* value as large as the observed. $\sigma$ represents the square root of the estimated random effect variance. The term ‘Block’ was dropped from the random effect specification, as it led to model overfitting.

| **Effect** | **Group** | **Term** | **Estimate** | **Std. error** | ***Z*** | ***p*** |
|---|---|---|---|---|---|---|
| fixed | NA | Intercept | 5.396 | 0.049 | 109.21 | <0.01 |
| fixed | NA | +N | -0.116 | 0.072 | -1.610 | 0.107 |
| fixed | NA | +P | -0.185 | 0.073 | -2.535 | 0.011 |
| fixed | NA | +N:+P | 0.178 | 0.104 | 1.710 | 0.087 |
| random | Plot | $\sigma$ | 0 | NA | NA | NA |

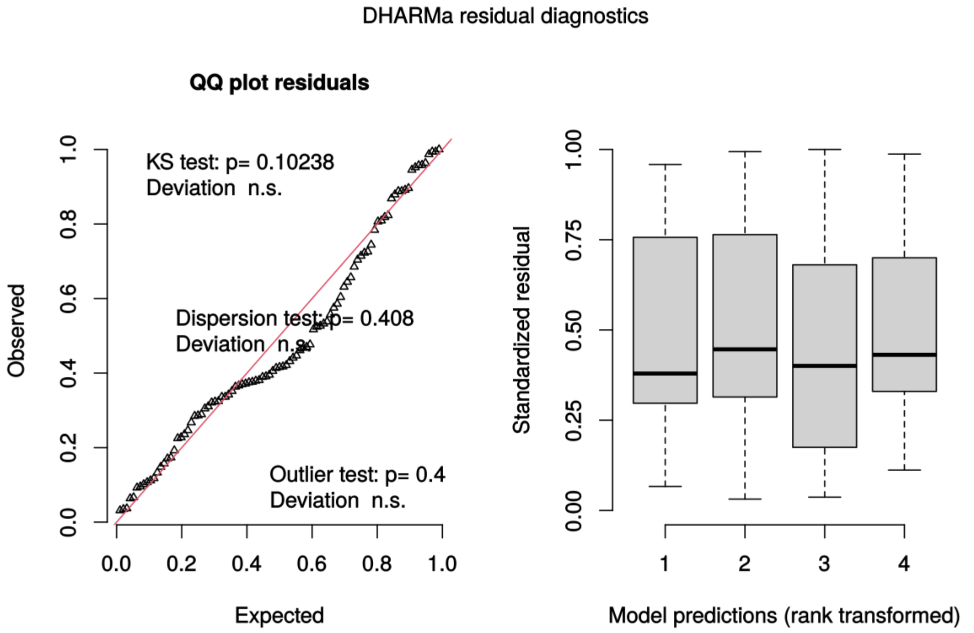

**Figure S3** Assumption validation of first glmm (Table S1) with the DHARMa R package. Both figures show that data fit well model assumptions, residuals are not overdispersed, and the negative binomial distribution was a fair choice to model the variance of richness estimates.

Non peer rev

**Table S2** Coefficients of the second generalized linear mixed effect model where ASV richness of focal fungal phyla was modelled as a function of fertilization treatments. Estimates along the row labelled as 'intercept' correspond to the mean ASV richness of Ascomycota in control plots. Estimates and standard errors are presented in the natural logarithm scale, as this was the specified link function for the negative binomial probability distribution. *Z* represents the Wald-*Z* statistic, while *p* represents the probability of observing a theoretical *Z* value as large as the observed. $\sigma$ represents the square root of the estimated variance for each component of the nested random term.

| **Effect** | **Group** | **Term** | **Estimate** | **Std. error** | ***Z*** | ***p*** |
|---|---|---|---|---|---|---|
| fixed | NA | Intercept | 4.995 | 0.069 | 72.338 | 0.000 |
| fixed | NA | +N | -0.137 | 0.098 | -1.404 | 0.160 |
| fixed | NA | +P | -0.148 | 0.099 | -1.499 | 0.134 |
| fixed | NA | Basidiomycota | -1.305 | 0.077 | -16.867 | 0.000 |
| fixed | NA | Mortierellomycota | -2.860 | 0.099 | -28.792 | 0.000 |
| fixed | NA | +N:+P | 0.190 | 0.139 | 1.365 | 0.172 |
| fixed | NA | +N:Basidiomycota | 0.069 | 0.110 | 0.628 | 0.530 |
| fixed | NA | +N:Mortierellomycota | -0.014 | 0.143 | -0.094 | 0.925 |
| fixed | NA | +P:Basidiomycota | 0.005 | 0.112 | 0.045 | 0.964 |
| fixed | NA | +P:Mortierellomycota | -0.045 | 0.146 | -0.307 | 0.759 |
| fixed | NA | +N:+P:Basidiomycota | 0.059 | 0.157 | 0.379 | 0.705 |
| fixed | NA | +N:+P:Mortierellomycota | 0.253 | 0.203 | 1.241 | 0.214 |
| random | Plot:Block | $\sigma$ | 0.093 | NA | NA | NA |
| random | Block | $\sigma$ | 0.000 | NA | NA | NA |

Non peer reviewed

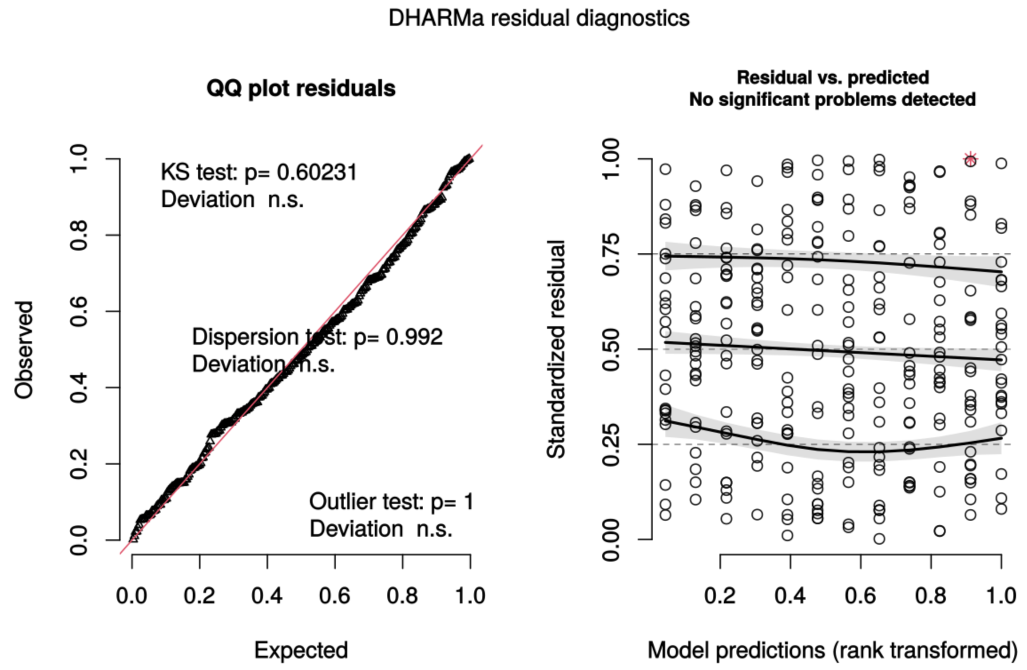

**Figure S4** Second glmm (Table S2) assumption validation with the DHARMa R package. Both figures show that data fit well model assumptions: residuals are not overdispersed, and the negative binomial distribution was a fair choice to model the variance of richness estimates.

Non peer re

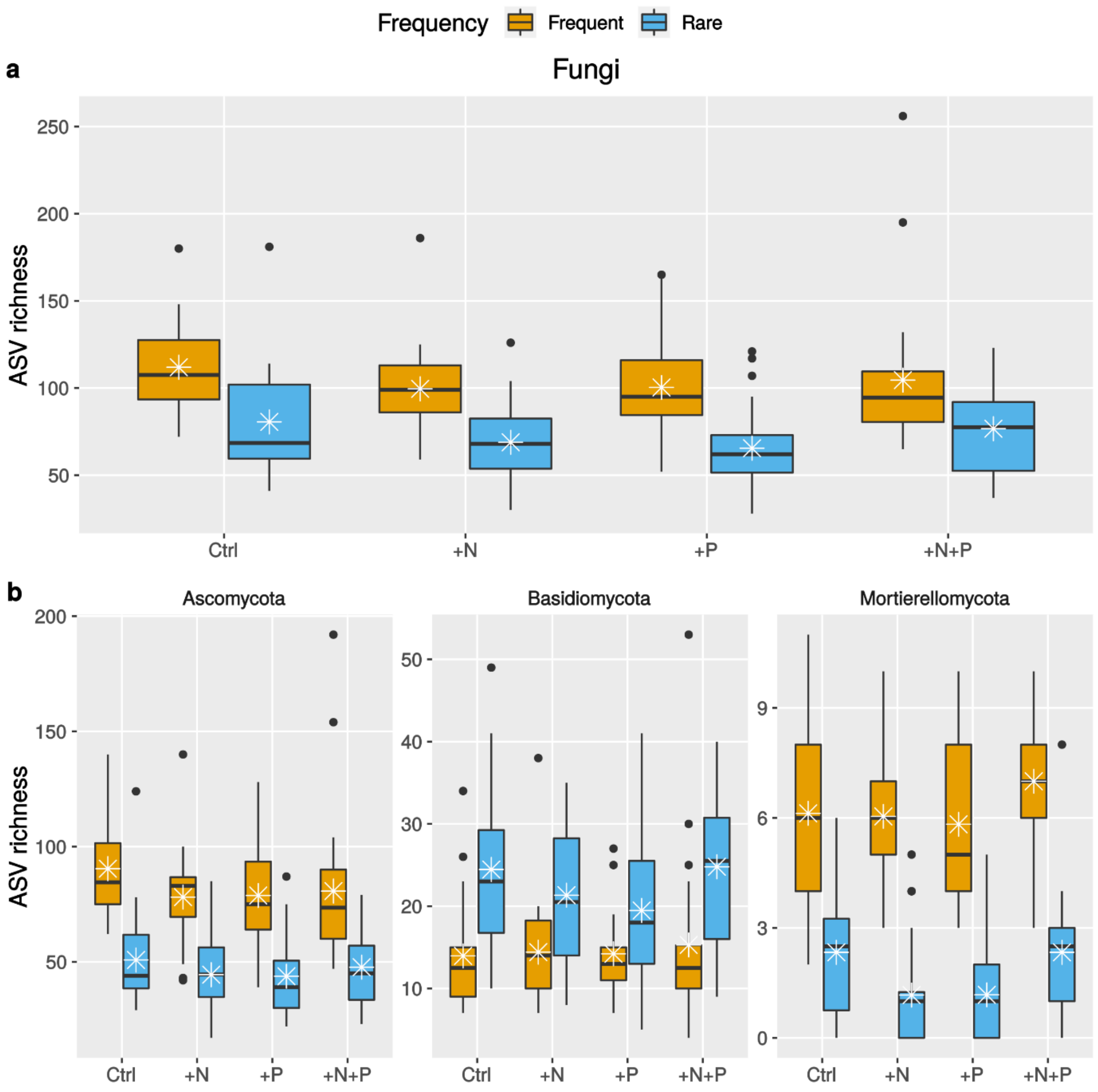

**Figure S5** Comparison of observed differences in ASV richness as a function of fertilization between frequent and rare variants (n=95). Variants which occurred in at least five samples or more were considered frequent. The horizontal bar in each box represents the median, while the white star represents the mean. Dots above each box represent outliers.

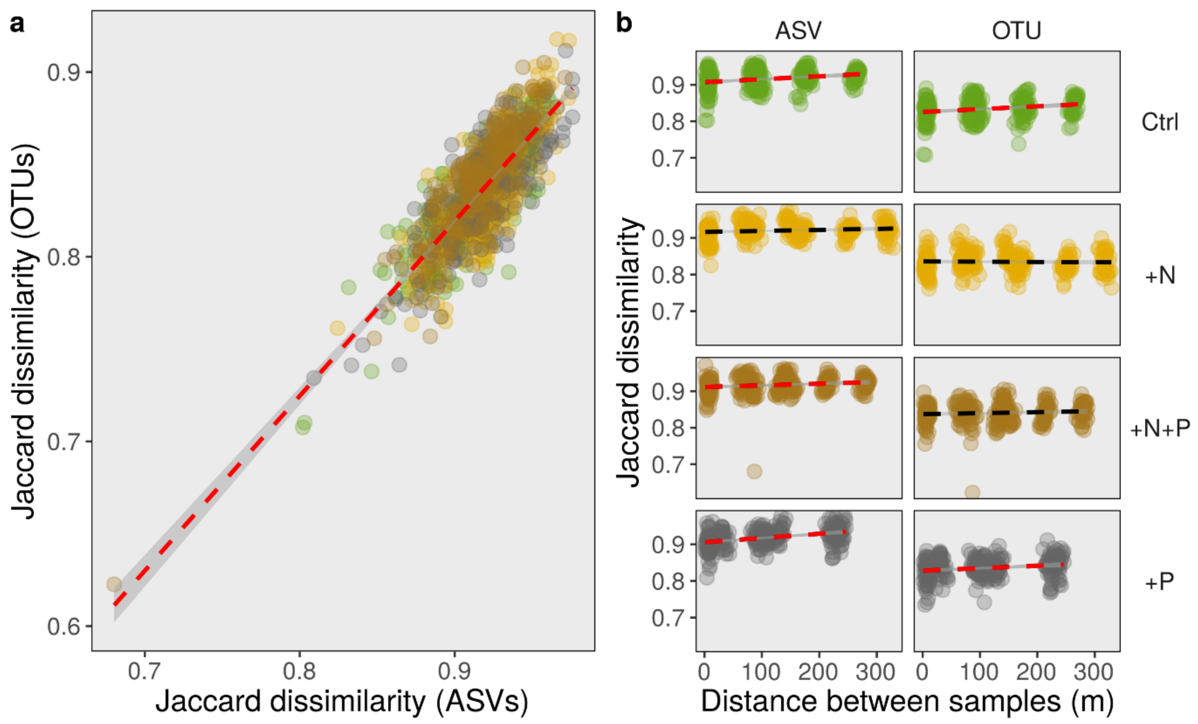

**Figure S6** Beta diversity of RAF communities. Beta diversity, defined as turnover of features across communities, is represented here by the Jaccard dissimilarity index. Communities composed by ASVs or by variants grouped within their corresponding species were used to visualize these relations. Panel **a** plots a correlation between beta diversity estimates of the finer levels of taxonomic resolution. Panel **b** plots the relation between beta diversity and Euclidean distance between samples. Red lines indicate the correlation was significant. Correlation parameters can be found in Table S3.

Non peer rev

**Table S3** Mantel correlation tests divided by operational taxonomic unit resolution and treatment. Jaccard dissimilarity matrices were correlated to Euclidean distance matrices. The default correlation type was employed (Pearson). Permutation test were free (n=999). OTUs are clustered at 97% similarity.

| **Taxonomic unit resolution** | **Treatment** | **Mantel $\rho$** | ***p*** |
|---|---|---|---|
| ASV | Control | 0.246 | 0.002 |
| ASV | +N | 0.129 | 0.07 |
| ASV | +N+P | 0.157 | 0.027 |
| ASV | +P | 0.355 | 0.001 |
| OTU | Control | 0.217 | 0.004 |
| OTU | +N | -0.019 | 0.554 |
| OTU | +N+P | 0.085 | 0.151 |
| OTU | +P | 0.188 | 0.027 |

Non peer reviewed

**Table S4** Tests of different neighborhood matrices to draw spatial eigenvectors. Residuals were extracted from multiple linear models where ASV tables were modelled as a function of spatial coordinates. Spatial coordinates were expressed in UTMs. Permutation tests were employed to select optimal neighborhood matrices. Optimal matrices maximize the proportion of variance explained (adj. $R^2$) by Moran eigenvector maps when significant positive correlations are found. Three graphical agglomerative criteria were tested: Gabriel graph (Gabriel), relative neighborhood (Relative) and minimum spanning tree (MST). Binary, linear and concave weighting parameters were also tested.

| | | **All ASVs** | | | **Frequent ASVs only** | |
|---|---|---|---|---|---|---|
| **Taxonomic level** | **Neighborhood criterion** | **adj. $R^2$** | ***p*** | | **adj. $R^2$** | ***p*** |
| Fungi | Gabriel - binary | 0.014 | 0.073 | | 0.031 | 0.020 |
| | Gabriel - linear | NA | NA | | 0.034 | 0.008 |
| | Relative - binary | 0.005 | 0.554 | | NA | NA |
| Ascomycota | Gabriel – binary | 0.018 | 0.047 | | 0.026 | 0.036 |
| | Gabriel – linear | 0.019 | 0.032 | | 0.032 | 0.008 |
| | Gabriel – concave down | 0.020 | 0.028 | | 0.027 | 0.039 |
| Basidiomycota | Gabriel - binary | 0.007 | 0.511 | | 0.012 | 0.588 |
| | Relative - binary | 0.000 | 0.853 | | 0.012 | 0.563 |
| | MST - binary | 0.014 | 0.219 | | 0.022 | 0.341 |
| Mortierellomycota | Gabriel - binary | 0.007 | 0.791 | | 0.018 | 0.663 |
| | Relative - binary | 0.010 | 0.728 | | 0.024 | 0.622 |
| | MST - binary | 0.005 | 0.780 | | 0.010 | 0.749 |

Non peer reviewed

**Table S5** Sensitivity tests of the effects of long term fertilization on the structure of root associated fungal (RAF) communities. Pseudo-*F* tests assessed the statistical significance of the proportion of compositional variance explained by fertilization treatments. Tests were repeated twice dividing the dataset in frequent and rare variants. Frequently observed (i.e. presence-absence matrices of those ASVs occurring in at least 5 samples) and rare variants were modelled as a function of fertilization factors. Only the best represented fungal clades recovered in this study were tested with the exception of Glomeromycota. Values of *p* were calculated with permutation tests (n = 9999).

| | | **Frequent ASVs only** | | | **Rare ASVs only** | |
|---|---|---|---|---|---|---|
| **Taxonomic level** | **Factor** | **Pseudo *F*** | ***p*** | | **Pseudo *F*** | ***p*** |
| Fungi | +N | 1.521 | 0.001 | | 1.007 | 0.243 |
| | +P | 1.377 | 0.005 | | 1.145 | 0.000 |
| | +N+P | 1.197 | 0.031 | | 1.023 | 0.301 |
| Ascomycota | +N | 1.752 | 0.000 | | 1.078 | 0.062 |
| | +P | 1.553 | 0.001 | | 1.171 | 0.007 |
| | +N+P | 1.188 | 0.038 | | 1.061 | 0.112 |
| Basidiomycota | +N | 1.154 | 0.217 | | 1.004 | 0.296 |
| | +P | 1.275 | 0.058 | | 1.134 | 0.002 |
| | +N+P | 1.376 | 0.026 | | 1.014 | 0.396 |
| Mortierellomycota | +N | 1.332 | 0.157 | | 0.882 | 0.535 |
| | +P | 1.516 | 0.088 | | 1.152 | 0.154 |
| | +N+P | 1.488 | 0.104 | | 1.058 | 0.346 |

Non peer reviewed

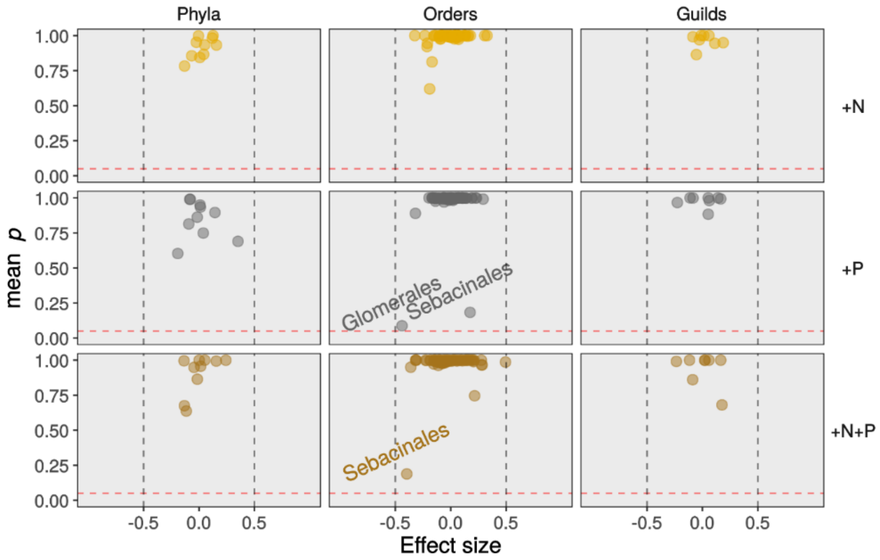

**Figure S7** Graphical illustration of the results of univariate tests of the effect of fertilization treatments on the relative read abundance of taxonomic clades and trophic guilds. Data was modelled by generalized linear models (glm). Abundance was calculated by aggregating ASV read counts according to their phylum (n=10), order (n=91) or guild (n=8) classification. Read counts were then transformed to $log_2$ ratios using the center log ratio procedure. Plots in the grid show the relation between the standardized effect size (median difference in relation to control / maximum difference within treatment) and the mean *p* value. Expected *p* values were corrected by the Benjamini-Hochberg procedure in order to minimize the probability of false positives due to multiple testing. The dashed horizontal line references a *p* value of 0.05.

Non peer reviewed
